# An ACAT inhibitor regulates SARS-CoV-2 replication and antiviral T cell activity

**DOI:** 10.1101/2022.04.12.487988

**Authors:** Nathalie M Schmidt, Peter AC Wing, Rory Peters, Rachel Brown, Hao Wang, Leo Swadling, COVIDsortium Investigators, Joseph Newman, Nazia Thakur, Kaho Shionoya, Sophie B Morgan, Timothy SC Hinks, Koichi Watashi, Dalan Bailey, Scott B Hansen, Mala K Maini, Jane A McKeating

**Author notes:** Joint first authors (Nathalie M Schmidt, Peter AC Wing). Joint corresponding authors. listed in appendix (COVIDsortium Investigators).

## Abstract

The severity of disease following infection with SARS-CoV-2 is determined by viral replication kinetics and host immunity, with early T cell responses and/or suppression of viraemia driving a favourable outcome. Recent studies have uncovered a role for cholesterol metabolism in the SARS-CoV-2 life cycle and in T cell function. Here we show that blockade of the enzyme Acyl-CoA:cholesterol acyltransferase (ACAT) with Avasimibe inhibits SARS-CoV-2 entry and fusion independent of transmembrane protease serine 2 expression in multiple cell types. We also demonstrate a role for ACAT in regulating SARS-CoV-2 RNA replication in primary bronchial epithelial cells. Furthermore, Avasimibe boosts the expansion of functional SARS-CoV-2-specific T cells from the blood of patients sampled in the acute phase of infection. Thus, re-purposing of available ACAT inhibitors provides a compelling therapeutic strategy for the treatment of COVID-19 to achieve both antiviral and immunomodulatory effects.

## Introduction

SARS-CoV-2 is a global health issue associated with over 400 million infections and 6 million deaths (WHO, 2022). Preventive vaccines have reduced morbidity and mortality (Gupta et al., 2021b; Sheikh et al., 2021); however, therapeutic strategies for unvaccinated subjects or those with breakthrough infections are needed. Several direct-acting antiviral drugs are now licensed for the treatment of SARS-CoV-2 infection (Molnupiravir and Nirmatrelvir), whilst other approaches boost host defences, for example, supplementing Type I interferon or neutralising antibodies (NIH, 2022). Accumulating data support an essential role for SARS-CoV-2-specific T cells in the early control of viraemia associated with mild, asymptomatic or even abortive infection (Moderbacher et al., 2020; Moss, 2022; Swadling et al., 2021; Tan et al., 2021). In contrast, T cells in patients with severe disease express exhaustion markers like programmed death-1 (PD-1) (Chen and Wherry, 2020), suggesting that approaches to restore T cell functionality may be beneficial. To the best of our knowledge there are currently no agents that show both direct antiviral and immune boosting activity against SARS-CoV-2 infection.

Metabolic syndrome and hyperlipidaemia have been associated with a poorer outcome of SARS-CoV-2 infection and cholesterol-lowering HMG-CoA-reductase inhibitors (statins) may improve COVID-19 survival, highlighting the potential of targeting cholesterol metabolism as a treatment strategy (Bergqvist et al., 2021; Gupta et al., 2021a; Schmidt et al., 2020). Cholesterol is a key component of cellular membrane lipids regulating curvature, fluidity and the formation of microdomains or lipid rafts in the plasma membrane that are sites of receptor signalling (Ikonen, 2008). Cholesterol homeostasis is integral to many steps in the life cycle of a wide range of viruses, including entry, replication, assembly and egress (Glitscher and Hildt, 2021) and recent studies have identified a role in SARS-CoV-2 particle infectivity, syncytia formation and genome replication (Daniloski et al., 2021; Palacios-Rápalo et al., 2021; Sanders et al., 2021). In immune cells, cholesterol availability, uptake and utilization are linked to immune function and shape antiviral responses (Kidani et al., 2013; Schmidt et al., 2020; Spann and Glass, 2013).

Acyl-CoA:cholesterol acyltransferase (ACAT, also known as sterol O-acyltransferase, SOAT) esterifies free cholesterol; pharmacological inhibition of ACAT reduced hepatitis B and C virus replication (Hu et al., 2017; Schmidt et al., 2021), whilst enhancing antiviral and anti-tumour T cell responses (Schmidt et al., 2021; Yang et al., 2016). We previously reported that ACAT inhibition induced metabolic reprogramming to preferentially boost the exhausted T cell response that is characteristic of chronic hepatitis B virus infection and hepatocellular carcinoma (Schmidt et al., 2021). We found that cholesterol-rich microdomains required for T cell synapse formation and antigen recognition were reduced in exhausted T cells expressing high levels of PD-1 (PD-1^hi^) and ACAT inhibition restored these properties, suggesting it may provide beneficial effects on the activated PD-1^hi^ antiviral T cells in acute SARS-CoV-2 infection. Thus, we hypothesized that modulation of cholesterol metabolism by ACAT inhibitors such as Avasimibe (AVS) would inhibit SARS-CoV-2 replication and boost virus-specific T cells to control infection.

## Results

### Avasimibe blocks SARS-CoV-2 pseudoparticle entry

SARS-CoV-2 infection is mediated by Spike protein binding to angiotensin-converting enzyme (ACE2) that enables cleavage by the transmembrane protease serine 2 (TMPRSS2), triggering fusion of viral and host membranes at the cell surface (Hoffmann et al., 2020; Wan et al., 2020). However, SARS-CoV-2 can infect cells lacking TMPRSS2 where particles enter by ACE2-dependent endocytosis with fusion occurring in endosomal vesicles (Jackson et al., 2022). To assess whether ACAT inhibition with AVS can regulate plasma membrane or endosomal viral fusion we used lentiviral pseudoparticles bearing the SARS-CoV-2 Spike (Victoria 01/20 strain) to study cell entry in VeroE6 cells that lack TMPRSS2 or cells engineered to over-express this serine protease. Pre-treatment of cells with AVS reduced pseudoparticle infection of both VeroE6 and VeroE6-TMPRSS2 cells (**Fig.1a**). AVS has been reported to alter plasma membrane cholesterol levels, showing a reduction in hepatoma cells (Jiang et al., 2019) and an increase in CD8^+^ T cells (Schmidt et al., 2021; Yang et al., 2016), suggesting cell-type differences. Cholesterol plasma membrane levels in the AVS treated VeroE6 cells showed a modest but significant increase in free cholesterol with no change in total levels, consistent with a redistribution from cholesteryl ester stored in lipid droplets to unesterified membrane cholesterol (**Supp.Fig.1a**). Membrane cholesterol can cluster in lipid rafts, cholesterol- and glycosphingolipid-rich microdomains that can be identified by fluorescent-labelled cholera toxin B (CTB) subunit binding to monosialotetrahexosylganglioside (GM1) and visualized by direct stochastical optical reconstruction microscopy (dSTORM). AVS increased the diameter of GM1-enriched domains in VeroE6 cells (**Supp.Fig.1b**), providing further evidence that AVS treatment increased plasma membrane cholesterol.

**Figure 1.**
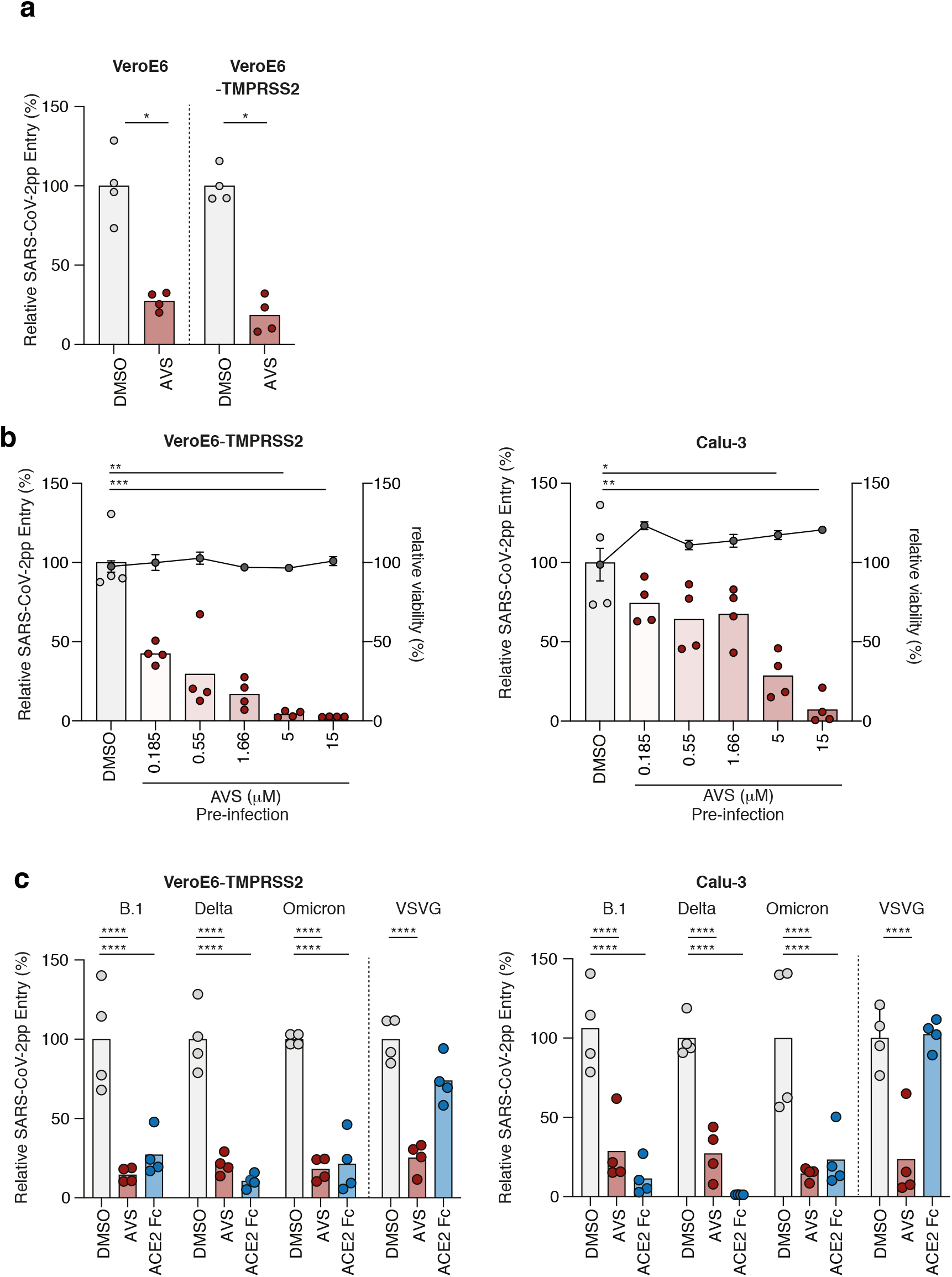
Avasimibe blocks SARS-CoV-2 pseudoparticle entry. **(a)** VeroE6 and VeroE6-TMPRSS2 cells were treated with 20μM of Avasimibe (AVS) for 24h prior to infection with lentiviral pseudoparticles (pp) bearing the SARS-CoV-2 spike protein (VIC 01/20) and luciferase activity measured 48h post-infection. **(b)** Vero-TMPRSS2 (left) and Calu-3 (right) cells were pre-treated for 24h with AVS (red) or DMSO (light grey) and infected with SARS-CoV-2pp (VIC 01/20). Luciferase activity and cell viability (dark grey) were measured 48h post-infection and data is representative of n=4 biological replicates. **(c)** Viral pp were generated bearing spike proteins from B.1, Delta, and Omicron variants of concern or VSV-G and used to infect VeroE6-TMPRSS2 (left) or Calu-3 (right) cells pre-treated with 20μM of AVS. As a control to evaluate ACE2-dependency of infection all pp were incubated with 1μg/ml of ACE2-Fc prior to infecting target cells. All data are normalized to mean of DMSO and P values determined by ANOVA (Kruskal Wallis).

To extend our observations to a lung epithelial cell line, we selected Calu-3 cells that express endogenous ACE2/TMPRSS2. We observed a dose-dependent inhibition of SARS-CoV-2 pseudoparticle infection with an IC_50_ of 1.77μM of AVS compared to 0.23μM for Vero-TMPRSS2 cells and no detectable effect on cell viability (**Fig.1b**). The emergence of SARS-CoV-2 variants of concern (VOC) with altered Spike proteins such as Delta and Omicron that can partially evade vaccine protection prompted us to evaluate their sensitivity to ACAT inhibition. AVS inhibited the infection of both VeroE6-TMPRSS2 and Calu-3 by pseudoparticles expressing B.1 (D614G), Delta and Omicron Spike proteins (**Fig.1c**). As a control we showed that all SARS-CoV-2 pseudoparticles were neutralized with a saturating dose of ACE2-Fc, confirming ACE2 dependent entry. To evaluate whether this antiviral activity was dependent on endocytic trafficking we infected cells with pseudoparticles bearing Vesicular Stomatitis Virus G glycoprotein (VSV-G) that internalizes via clatherin-dependent endocytosis and fuses with endosomal membranes (Podbilewicz, 2014). AVS reduced VSV-G pseudoparticle infection of both cell lines (**Fig.1c**), reinforcing a role for AVS in perturbing endocytic trafficking pathways.

### Avasimibe blocks SARS-CoV-2 entry and replication

To determine whether our observations with lentiviral pseudoparticles translate to authentic viral replication, we pre-treated Calu-3 cells with AVS, infected with SARS-CoV-2 (Victoria 01/20 strain) and showed a significant reduction in intracellular viral RNA (**Fig.2a**). To examine whether ACAT regulates post-entry steps we treated Calu-3 or Vero-TMPRSS2 cells with AVS post-infection and showed a reduction in viral RNA in both cell types, with IC_50_ values of 5.99μM or 1.67μM, respectively (**Fig.2b**). Infected cells were treated with the nucleoside analogue remdesivir and we noted comparable inhibition of infection to AVS. Finally, we assessed the effect of AVS on SARS-CoV-2 infection of human primary bronchial epithelial cells (PBEC) grown at air-liquid-interface to provide a more physiological model of infection. Treatment with AVS pre- or post-infection reduced viral RNA and infectious virus shed from the apical surface of the cultures (**Fig.2c**). Taken together, ACAT inhibition has a direct antiviral effect against SARS-CoV-2 entry and RNA replication.

**Figure 2.**
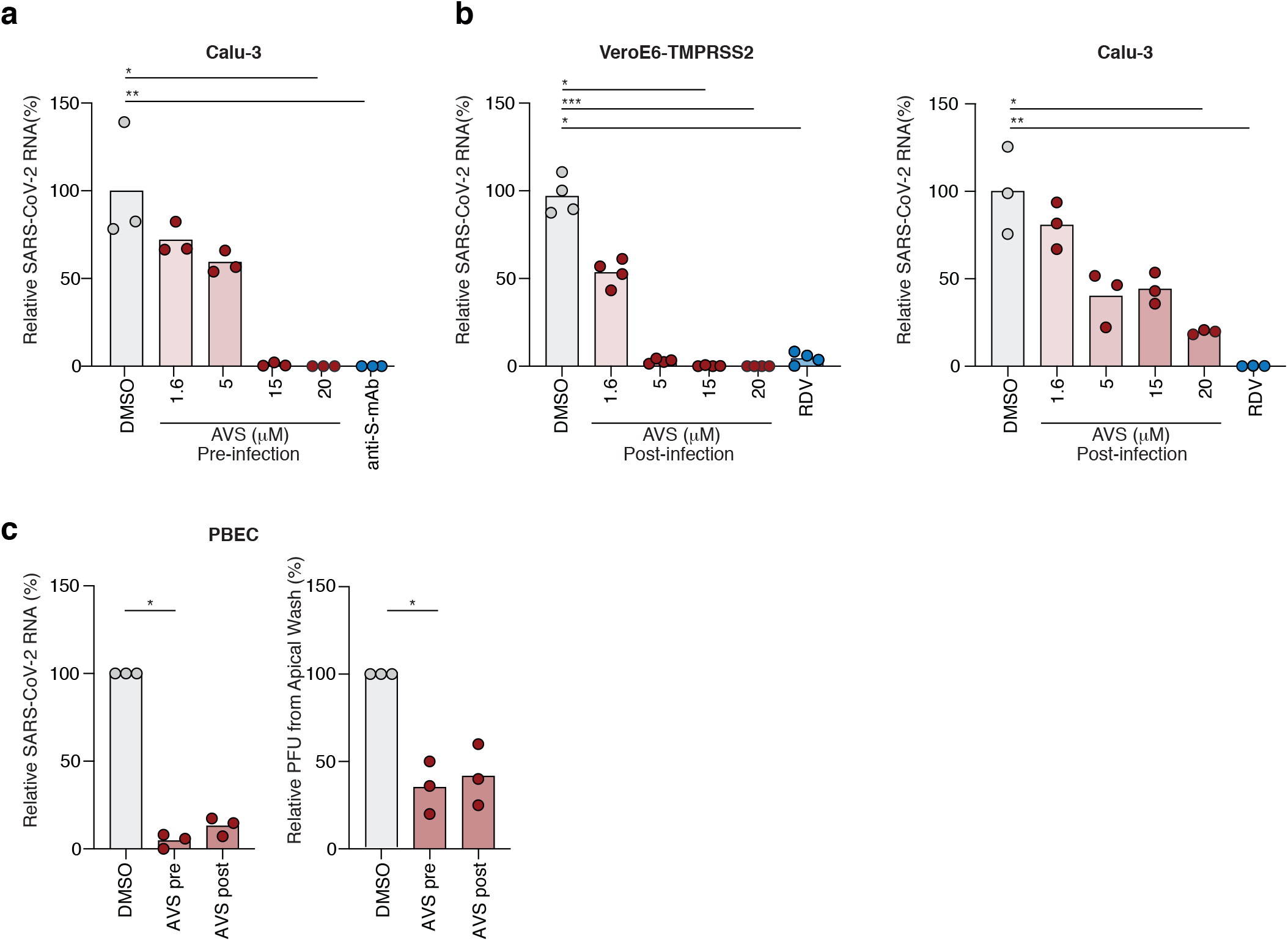
Avasimibe blocks SARS-CoV-2 entry and replication. **(a)** Calu-3 cells were pre-treated with AVS for 24h prior to infection with SARS-CoV-2 (VIC 01/20) at an MOI of 0.01. Cells were harvested 24h post infection and intracellular viral RNA quantified by qPCR. Data is representative of n=3-4 biological replicates. **(b)** VeroE6-TMPRSS2 (left) and Calu-3 (right) cells were infected with SARS-CoV-2 (MOI 0.01) for 2h, the inoculum was removed, and the cells treated with AVS. Cells were harvested 24h post infection and intracellular viral RNA quantified by qPCR. Data is representative of n=4 biological replicates. **(c)** Primary bronchial epithelial cells (PBEC) grown to air-liquid-interface were treated with 20μM of AVS either 24h pre- or 2h post infection of the apical surface with SARS-CoV-2 (MOI 0.1). Cultures were harvested 24h post infection and viral RNA quantified by qPCR and infectious virus shed from the apical surface by viral plaque assay. Data is representative of n=3 donors. All data are normalized to mean of DMSO and P values determined by ANOVA (Kruskal Wallis).

### Impact of Avasimibe on SARS-CoV-2-specific T cells

Next, we examined the effect of AVS on SARS-CoV-2-specific T cell activity. PBMC isolated from the blood of unvaccinated patients hospitalised during the first pandemic wave in the UK (March-July 2020) were collected during PCR-confirmed SARS-CoV-2 infection (information about patient cohort in methods). PBMC were stimulated with peptide pools derived from virus encoded Spike and Membrane (Mem) proteins in the presence or absence of AVS. After short-term 8d culture, we measured key antiviral effector functions of antigen-specific CD4^+^ and CD8^+^ T cells by multiparameter flow cytometry (gating strategy **Supp.Fig.2a**). AVS increased the frequencies of CD4^+^ T cells producing the antiviral cytokines IFNγ, TNF (or both) and MIP1β in response to either spike or membrane peptides, boosting responses in some patients and inducing *de novo* responses in others (**Fig.3a-c, Supp.Fig.2b,c**). The response to AVS was heterogeneous, showing a 50-fold increase in the magnitude of IFNγ-producing T cells in one patient and decreased cytokine production in a minority of patients, as previously reported for other *in vitro* and *in vivo* immunotherapeutic approaches (Bengsch et al., 2014; Maini and Pallett, 2018). A similar enhancement was seen for cytokine-producing CD8^+^ T cells in individual donors but was less consistent than for CD4^+^ T cells, resulting in no overall significant changes for CD8^+^ T cell responses across the cohort (**Supp.Fig.2d-f**). CD4^+^ T cells provide help to activate and differentiate B cells, for example via the interaction of CD40 and CD40L (CD154). AVS increased the SARS-CoV-2-specific expression of CD154 (CD40L) on CD4^+^ T cells, reflecting an enhanced capacity to co-stimulate CD40 and to activate B cells (**Fig.3d**). Consistent with the expansion in frequencies of functional responses, AVS increased the proliferation of virus-specific CD4^+^ T cells (detected by CFSE dilution, **Fig.3e**). Immunomodulatory therapies for viral infections carry the risk of increasing bystander immune responses and cytotoxic tissue damage; however, we did not detect any significant increase of CD107a mobilization to the cell membrane of perforin-producing T cells, markers of degranulation and cytotoxicity respectively (**Supp.Fig.2g**). COVID-19 severity is associated with male sex (Scully et al., 2020) and increased age (Richardson et al., 2020). We noted that AVS enhancement of SARS-CoV-2-specific T cell responses was seen in both males and females and was independent of age (**Supp.Fig.2h**), showing the potential of this therapeutic approach for a variety of patients, including those at risk of severe infection.

**Figure 3.**
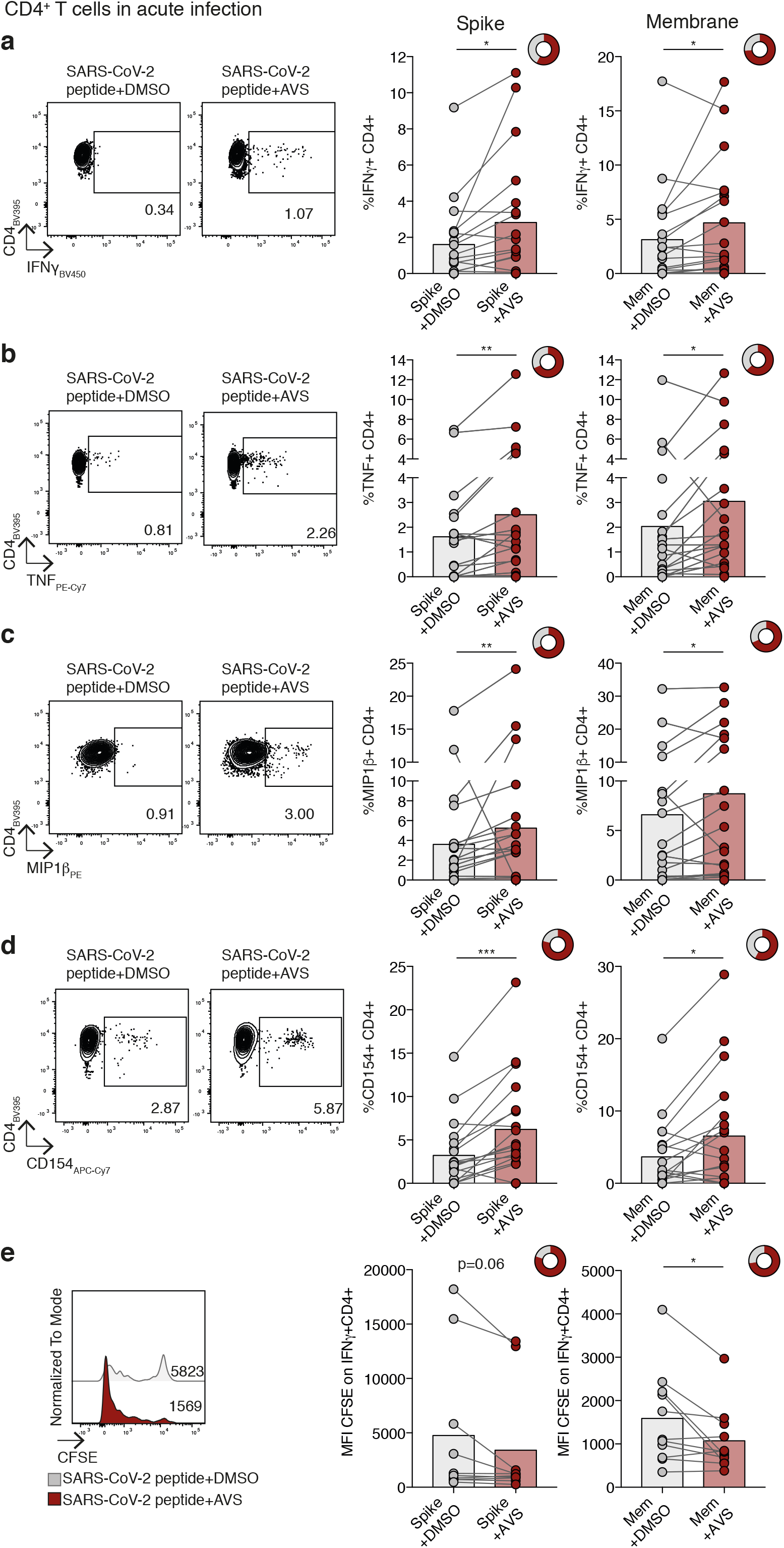
Impact of Avasimibe on SARS-CoV-2-specific CD4^+^ T cells in acute infection. **(a-e)** Human PBMC from donors with acute SARS-CoV-2 infection were stimulated with SARS-CoV-2 peptide pools (Spike and Membrane, Mem) and treated with Avasimibe (AVS) or DMSO for 8d. SARS-CoV-2-specific cytokine production by CD4^+^ T cells was detected via flow cytometry. The cytokine production/CD154 expression in wells without peptide stimulation was subtracted to determine SARS-CoV-2-specific cytokine production/CD154 expression in summary data. Example plots and summary data for SARS-CoV-2 specific IFNγ(**a**), TNF (**b**), MIP1β(**c**) production and CD154 expression (**d**) by CD4^+^ T cells (n=19). (**e**) Assessment of SARS-CoV-2-specific proliferation determined by CFSE dilution gated on IFNγ^+^ CD4^+^ T cells (Spike n=10; Mem n=11). Bars mean. Doughnut charts indicate fraction of donors with response to AVS (red). Response defined as *de novo* or increased cytokine production/CD154 expression. P values determined by Wilcoxon matched-pairs signed rank test.

To ascertain whether AVS only boosts virus-specific effector and early memory T cells during or shortly after infection but not memory T cells, we recruited a second cohort of unvaccinated donors 6 months after SARS-CoV-2 infection (memory cohort, see methods section). AVS had no consistent effect on SARS-CoV-2-specific memory CD4^+^ or CD8^+^ T cell responses 6 months post-infection (**Supp.Fig.3a-d**). This is in line with our previous findings showing that ACAT inhibition preferentially rescues PD-1^hi^ T cells and not memory responses to cytomegalovirus (Schmidt et al., 2021). AVS has shown a good safety profile in phase III atherosclerosis studies (Llaverías et al., 2003) and has not been associated with autoimmune responses in murine models (Yang et al., 2016). In line with this, we did not detect any non-specific increase in cytokine production when T cells from the acute cohort were treated with AVS without viral peptides (**Supp.Fig.3e**). Thus, our data support AVS selectively expanding acutely activated SARS-CoV-2-specific T cells, without affecting memory or non-activated T cells.

## Discussion

This study raises a number of areas for future investigation. AVS inhibition of SARS-CoV-2 fusion in VeroE6 and VeroE6-TMPRSS2 is consistent with ACAT regulating cholesterol levels at both the cell surface and within endosomes, highlighting the need to better understand the role of cholesterol in endosomal pathways that are essential in virus internalization and egress (Glitscher and Hildt, 2021). Our observation that AVS inhibited VSV-G pseudoparticle entry suggests a potential role in regulating the entry of other viruses that would be worth investigating. Cholesterol 25-hydroxylase catalyzes the formation of 25-hydroxycholesterol (25HC) from cholesterol and leads to a redistribution of cholesterol limiting the entry of a range of enveloped viruses (Schoggins, 2019) including SARS-CoV-2 (Wang et al., 2020; Zang et al., 2020; Zu et al., 2020). Wang et al reported that 25HC activated ACAT and suggested this as a mechanism to explain 25HC inhibition of SARS-CoV-2 entry. The authors showed that inhibition of ACAT with SZ58-035 partially reversed the antiviral activity of 25HC in Calu-3 cells; however, they observed a negligible effect on basal plasma membrane cholesterol levels or on SARS-CoV-2 pseudoparticle entry. This contrasts with our results and may reflect variable efficacy of SZ58-035 and AVS to modulate cholesterol levels. Our observation that AVS inhibits SARS-CoV-2 pseudoparticle infection of a range of cell lines and primary epithelial cells shows its robust antiviral activity.

We focused on T cells specific for two of the key structural proteins targeted in acute infection (Peng et al., 2020) and further studies to assess the effect of AVS on other T cell specificities including those against non-structural viral proteins associated with abortive infection would be of interest (Swadling et al., 2021). The potential for AVS to boost acutely activated CD4^+^ T effector and helper function even in the elderly, suggests they could be tested for their capacity to adjuvant sub-optimal vaccine responses in this vulnerable group (Collier et al., 2021) or others with waning immunity. The lack of T cell boosting in the memory phase is in line with our previous findings (Schmidt et al., 2021) but conceivably could also be related to the younger age of this cohort.

We have shown increased antiviral activity following treatment of circulating T cells; however immune responses at the site of disease, the lung and upper respiratory tract, are shaped by the local microenvironment and nutrient availability. The lung is enriched in cholesterol compared to blood (Chamberlain, 1928) with cholesterol constituting the main neutral lipid in surfactant (Keating et al., 2007). We previously reported that ACAT inhibition is enhanced in the presence of high cholesterol. T cells isolated from cholesterol-rich liver and tumour tissues were boosted to a greater extent than those from the blood of the same donors (Schmidt et al., 2021); suggesting a similar enhancement may be seen following ACAT inhibition of SARS-CoV-2-specific T cells infiltrating the infected lung. Further studies to address the effect of AVS on other immune cell subsets associated with the inflammatory response in severe and long COVID-19 would also be of interest.

Urgent consideration should be given to trials testing the efficacy of re-purposing ACAT inhibitors like AVS, an oral agent that has been shown to have a good safely profile. We show it has the capacity to exert a unique dual effect, directly inhibiting SARS-CoV-2 entry and RNA replication as well as boosting the acute T cell response that can aid viral elimination and provide protection against re-infection.

## Methods

### Ethics

The COVIDsortium cohort was approved by the ethical committee of UK National Research Ethics Service (20/SC/0149) and registered at https://ClinicalTrials.gov (NCT04318314). The Royal Free Biobank (TapB) was approved by the Wales Research Ethics Committee (16/WA/0289; 21/WA/0388; project approval reference: NC2020.11). The PBEC study was reviewed by the Oxford Research Ethics Committee B (18/SC/0361). All study participants gave written informed consent prior to inclusion in the study and all storage of samples obtained complied with the Human Tissue Act 2004.

### Patient Cohort

Peripheral blood samples were taken from unvaccinated study participants during or after SARS-CoV-2 infection during the first pandemic wave of infections in the UK (March-July 2020).

The Acute Cohort was recruited from hospitalized patients at the Royal Free Hospital, London, and SARS-CoV-2 infection was confirmed by PCR (n=22; median age 82 years; 45% female, 55% male; 73% white, 4% black, 14% Asian, 9% other).

The Memory Cohort (COVIDsortium) was recruited from healthcare workers in London and SARS-CoV-2 infection was confirmed by PCR and/or serology. Samples were taken 5-6 months post infection (n=12; median age 44.5 years; 50% female, 50% male; 50% white, 8% black, 34% Asian, 8% other). More information about the cohort can be found in (Augusto et al., 2020).

### PBMC Isolation

For samples taken during acute SARS-CoV-2 infection, PBMC were isolated from EDTA blood using Histopaque-1077 (Sigma-Aldrich) density-gradient centrifugation in Leukosep tubes (Greiner Bio One) according to the manufacturer’s instructions. For COVIDsortium cohort, PBMC were isolated from heparinized blood samples using Pancoll (Pan Biotech) or Histopaque-1077 Hybri-Max (Sigma-Aldrich) density-gradient centrifugation in SepMate tubes (StemCell) according to the manufacturer’s instructions.

Isolated PBMC were cryopreserved in fetal bovine serum (FBS; Sigma-Aldrich) containing 10% dimethyl sulfoxide (DMSO; Sigma-Aldrich) and stored in liquid nitrogen prior to cell culture.

### Short-term cell culture

To examine SARS-CoV-2-specific T cell responses in the blood, PBMC were stimulated with 1μg/ml SARS-CoV-2 peptide pools (Membrane (Mem): 15mer peptides overlapping by 10aa, 43 peptides total; Spike S1: 18-20mer peptides, 18 peptides total. The full peptide sequences can be found in (Reynolds et al., 2020)) in cRPMI (RPMI 1640 (Thermo Fisher Scientific)+2% HEPES buffer solution, 0.5% sodium pyruvate, 0.1% 2-mercaptoethanol, MEM 1% non-essential and 2% essential amino acids; Gibco, and 100U/ml penicillin/streptomycin; life technologies)+10% FBS+20U/ml recombinant human IL-2 (PeproTech)+ 5μg/ml anti-CD28 (Invitrogen). PBMC were expanded at 37°C for 8d±0.5μM of the ACAT inhibitor Avasimibe (AVS; Selleckchem) or equivalent concentration of DMSO replenished every 2d. On d7, PBMC were restimulated with 1μg/mL peptide + anti-CD28 in the presence of 1μg/ml Brefeldin A (Sigma-Aldrich) for 16h at 37°C, followed by antibody staining and flow cytometric analysis. All experiments were performed in duplicates and combined prior to restimulation. Post-culture viability of PBMC was confirmed and samples with <50% viable cells were excluded from further analysis. The cytokine production/CD154 expression in wells without peptide stimulation was subtracted to determine peptide-specific cytokine production in all summary data. A SARS-CoV-2-specific response was defined as a minimum of 10 cells in the positive fraction. For evaluation of cell proliferation, PBMCs were labelled with 1μM CFDA-SE (Thermo Fisher) prior to the start of culture.

### Surface and intracellular staining

For flow cytometry, cells were stained with saturating concentrations of surface antibodies and a fixable viability dye diluted in 1:1 PBS (Invitrogen): Brilliant Violet Buffer (BD Biosciences). Following surface staining, cells were fixed and permeabilized with cytofix/cytoperm (BD Biosciences) followed by an intracellular staining with antibodies in saturating concentrations diluted in a 0.1% saponin-based buffer (Sigma-Aldrich). Full details on fluorescent monoclonal antibodies can be found in Supplementary Table 1. All samples were acquired on a BD Biosciences Fortessa-X20 or Fortessa and analysed using FlowJo v.10 (BD Biosciences).

### Human PBEC

Biopsies were obtained using flexible fibreoptic bronchoscopy from healthy control volunteers under light sedation with fentanyl and midazolam. Airway epithelial cells were taken using 2mm diameter cytology brushes from 3^rd^ to 5^th^ order bronchi and cultured in Airway Epithelial Cell medium (PromoCell) in submerged culture.

### SARS-CoV-2 pseudoparticle genesis and infection

Lentiviral pseudoparticles were generated by transfecting 293T cells with p8.91 (Gag-pol), pCSFW (luciferase reporter) and a codon optimised expression construct pcDNA3.1-SARS-CoV-2-Spike, as previously described (Thompson et al., 2020). Delta and Omicron Spike expression plasmids were provided by G2P-UK National Virology consortium. Supernatants containing viral pseudotypes were harvested at 48h and 7 h post-transfection. Viral titres were determined by infecting Calu-3 cells with a serial dilution of virus and 48h later measuring cellular luciferase. As a control for non-specific lentivirus uptake, stocks were generated with no envelope glycoprotein (No Env). This control was included in all experiments and luciferase values obtained subtracted from values acquired with the SARS-CoV-2pp. As an additional control pseudotypes were incubated with anti-S mAb F1-3A (1μg/mL) or ACE2-Fc (1μg/mL) for 30min prior to infection.

### SARS-CoV-2 propagation and infection

Naïve VeroE6 cells were infected with SARS-CoV-2 at an MOI of 0.003 and incubated for 48-72h until visible cytopathic effect was observed. At this point, cultures were harvested, clarified by centrifugation to remove residual cell debris and stored at −80°C. Viral titre was determined by plaque assay. Briefly, VeroE6 cells were inoculated with serial dilutions of SARS-CoV-2 viral stocks for 2h followed by addition of a semi-solid overlay consisting of 1.5% carboxymethyl cellulose (Sigma-Aldrich). Cells were incubated for 72h, visible plaques enumerated by fixing cells using amido black stain and plaque-forming units (PFU) per mL calculated. For infection of Calu-3 cells with SARS-CoV-2, cells were plated 24h before infection with the stated MOI. Cells were inoculated for 2h after which the residual inoculum was removed with three PBS washes. Unless otherwise stated, infected cells were maintained for 24h before harvesting for downstream applications.

### qPCR quantification of viral RNA

Total cellular RNA was extracted using the RNeasy kit (Qiagen) according to manufacturer’s instructions. For quantification of viral or cellular RNA, equal amounts of RNA, as determined by nanodrop, were used in a one-step RT-qPCR using the Takyon-One Step RT probe mastermix (Eurogentec) and run on a Roche Light Cycler 96. For quantification of viral copy numbers, qPCR runs contained serial dilutions of viral RNA standards. Total SARS-CoV-2 RNA was quantified using: 2019-nCoV_N1-F: 5’-GAC CCC AAA ATC AGC GAA AT-3’, 2019-nCoV_N1-R: 5’-TCT GGT TAC TGC CAG TTG AA TCT G-3’, 2019-nCoV_N1-Probe: 5’-FAM-ACC CCG CAT TAC GTT TGG TGG ACC-BHQ1-3’.

### Cholesterol measurement

To measure the relative changes in plasma membrane cholesterol after treatment with AVS, we developed an Amplex Red-based cholesterol detection assay. Briefly, VeroE6 cells were seeded into 96-well flat culture plates with transparent-bottom to reach confluency (~ 5 × 10^4^ per well). Cells were incubated with fresh EMEM+10%FBS for 1h followed by 1h of incubation in 100μL EMEM+10%FBS with 5μM AVS or equivalent concentrations of DMSO. After washing with 200μL PBS, cholesterol assay reactions were promptly begun by adding 100μL of working solution containing 50μM Amplex red, 1U/mL horseradish peroxidase, 2U/mL cholesterol oxidase and 2U/mL cholesterol esterase in PBS. Relative cholesterol concentration and the background (no cells) was determined in triplicates for each sample by measuring fluorescence activity with a fluorescence microplate reader (Tecan Infinite 200 PRO, reading from bottom) for 2h at 37°C with excitation wavelength of 530nm and an emission wavelength of 585nm. Subsequently, cholesterol level was normalized to the control activity after subtracting the background.

### dSTORM imaging of GM1

VeroE6 cells were grown to 30% confluence in EMEM+10%FBS. Cells were incubated with fresh EMEM+10%FBS for 1h followed by 1h of incubation in 100μL EMEM+10%FBS with 5μM AVS or equivalent concentrations of DMSO. Cells were rinsed with PBS and then fixed with 3% paraformaldehyde and 0.1% glutaraldehyde for 15min to fix both proteins and lipids. Fixative chemicals were reduced by incubating with 0.1% NaBH4 for 7min with shaking followed by three times 10min washes with PBS. Cells were permeabilized with 0.2% Triton X-100 for 15min and then blocked with a standard blocking buffer (10% bovine serum albumin (BSA) / 0.05% Triton in PBS) for 90min at room temperature. For labelling, cells were incubated with Alexa Fluor 647-CTB (Sigma-Aldrich) for 60min in 5% BSA / 0.05% Triton / PBS at room temperature followed by 5 washes with 1% BSA / 0.05% Triton / PBS for 15min each. Cells were then washed with PBS for 5min. Cell labelling and washing steps were performed while shaking. Labelled cells were then post-fixed with fixing solution, as above, for 10min without shaking followed by three 5min washes with PBS and two 3min washes with deionized distilled water.

Images were recorded with a Bruker Vutara 352 with a 60X Olympus Silicone objective. Frames with an exposure time of 20ms were collected for each acquisition. Excitation of the Alexa Fluor 647 dye was achieved using 640nm lasers and Cy3B was achieved using 561nm lasers. Laser power was set to provide isolated blinking of individual fluorophores. Cells were imaged in a photo-switching buffer comprising of 1%β-mercaptoethanol (Sigma-Aldrich), oxygen scavengers (glucose oxidase and catalase; (Sigma-Aldrich) in 50mM Tris (Affymetrix) + 10mM NaCl (Sigma-Aldrich) + 10% glucose (Sigma) at pH 8.0. Axial sample drift was corrected during acquisition through the Vutara 352’s vFocus system. Images were constructed using the default modules in the Zen software. Each detected event was fitted to a 2D Gaussian distribution to determine the centre of each point spread function plus the localization precision. The Zen software also has many rendering options including removing localization errors and outliers based on brightness and size of fluorescent signals. Pair correlation and cluster analysis was performed using the Statistical Analysis package in the Vutara SRX software. Pair Correlation analysis is a statistical method used to determine the strength of correlation between two objects by counting the number of points of probe 2 within a certain donut-radius of each point of probe 1. This allows for localization to be determined without overlapping pixels as done in traditional diffraction-limited microscopy. Cluster size estimation and cluster density were calculated through cluster analysis by measuring the length and density of the clusters comprising of more than 10 particles with a maximum particle distance of 0.1μm.

### Statistical analysis

Statistical analyses were performed with Prism 7.0 (GraphPad) as indicated in figure legends (Wilcoxon matched-pairs signed-rank test, Mann–Whitney test, Spearman correlation, Kruskall Wallis, unpaired t test) with significant differences marked on all figures. In experiments with a sample size >100 normality was assessed using a D’Agostino-Pearson omnibus normality test. All tests were performed as two-tailed tests, and for all tests, significance levels were defined as not significant (ns) *P*≥0.05; \**P*<0.05; \*\**P*<0.01; \*\*\**P*<0.001; \*\*\*\**P*<0.0001.

## Acknowledgements

We are grateful to all volunteers participating in this study and to the invaluable help of all clinical staff in sample acquisition. We thank Antonio Bertoletti for providing the SARS-CoV-2 peptide pools, and Hans Stauss, Amir Gander and other TapB staff, Dr Liã Arruda and members of the UCL Centre for Clinical Microbiology for providing the PBMC from acute SARS-CoV-2 infection. We acknowledge the G2P-UK National Virology consortium funded by MRC/UKRI (MR/W005611/1) and the Barclay Lab at Imperial College for providing the Omicron and Delta spike plasmids, Tiong Kit Tan and Alain Townsend for providing anti-S mAb and ACE2-Fc.

## Author contributions

NMS, PACW, MKM and JAM conceived the project; NMS, PACW, KW, MKM and JAM designed experiments; NMS, PACW, RP, HW, KS, KW generated data; NMS, PACW, SBH, MKM and JAM analysed and interpreted data; LS, RB, DB, JN, NT, SBM, TSCH and COVIDsortium provided essential reagents, patient samples and/or clinical data. NMS, PW, MKM and JAM prepared the manuscript. All authors provided critical review of the manuscript.

